# EXOSC10/Rrp6 is essential for the eight-cell embryo/morula transition

**DOI:** 10.1101/2021.10.06.463350

**Authors:** Fabrice G. Petit, Soazik P. Jamin, Pierre-Yves Kernanec, Emmanuelle Becker, Guillaume Halet, Michael Primig

## Abstract

The conserved 3’-5’ exoribonuclease EXOSC10/Rrp6 is required for gametogenesis, brain development, erythropoiesis and blood cell enhancer function. The human ortholog is essential for mitosis in cultured cancer cells. Little is known, however, about the role of *Exosc10* during embryo development and organogenesis. We generated an *Exosc10* knockout model and find that *Exosc10*^−/−^ mice show an embryonic lethal phenotype. We demonstrate that *Exosc10* maternal wild type mRNA is present in mutant oocytes and that the gene is expressed during all stages of early embryogenesis. Furthermore, we observe that EXOSC10 early on localizes to the periphery of nucleolus precursor bodies in blastomeres, which is in keeping with the protein’s role in rRNA processing and may indicate a function in the establishment of chromatin domains during initial stages of embryogenesis. Finally, we infer from genotyping data for embryonic days e7.5, e6.5 and e4.5 and embryos cultured *in vitro* that *Exosc10*^−/−^ mutants arrest at the eight-cell embryo/morula transition. Our results demonstrate a novel essential role for *Exosc10* during early embryogenesis, and they are consistent with earlier work showing that impaired ribosome biogenesis causes a developmental arrest at the morula stage.

## Introduction

Cell division requires a tightly orchestrated balance between synthesis, modification and degradation of RNAs. The conserved 3’-5’ exoribonuclease EXOSC10/Rrp6 contributes to this process in cooperation with the nuclear RNA exosome (Kilchert et al., 2016), regulatory proteins (Eberle et al., 2015; Wagschal et al., 2012), and perhaps alone *via* its ability to bind RNA (Wasmuth and Lima, 2017). Extensive work using model organisms such as yeast, fly and mouse has revealed numerous EXOSC10/Rrp6 substrates (Davidson et al., 2019; Gudipati et al., 2012; Kuai et al., 2005; Lardenois et al., 2011; Neil et al., 2009; Pefanis et al., 2015; Schneider et al., 2012; Xu et al., 2009), roles in rRNA processing (Briggs et al., 1998; Davidson et al., 2019), DNA double strand break repair (Domingo-Prim et al., 2019; Marin-Vicente et al., 2015), protein network interactors (Meldal et al., 2015), and the threedimensional structure of the enzyme (Makino et al., 2013; Midtgaard et al., 2006; Wasmuth et al., 2014). Targeted gene inactivation studies revealed important functions for mouse EXOSC10 in gametogenesis, erythropoiesis, B-cell enhancer function and forebrain development (Jamin et al., 2017; Mehta et al., 2021; Pefanis et al., 2015; Ulmke et al., 2021; Wu and Dean, 2020). Human EXOSC10 is important for cell division of cultured cancer cells (Blomen et al., 2015) and was associated with the polymyositis/scleroderma overlap syndrome (Bluthner and Bautz, 1992; Ge et al., 1992; Mahler and Raijmakers, 2007).

Mouse embryogenesis is initiated when the fertilized egg (zygote) progresses through three rounds of mitotic divisions to form an eight-cell embryo, which undergoes polarization and compaction at embryonic (e) day e2.5; reviewed in (Clift and Schuh, 2013). The embryo then develops into the morula, followed by the blastocyst, which contains a cavity (blastocoel) and the inner cells that differentiate into the primitive endoderm and the epiblast, thereby establishing, together with the trophectoderm, the first three cell lineages. These early developmental steps culminate in the implantation of the embryo at e4.5 (Chazaud and Yamanaka, 2016; Zernicka-Goetz et al., 2009). Initial stages of embryo development depend upon maternal transcripts that are gradually replaced during the maternal-to-zygotic transition (MZT) stage after zygotic genome activation (ZGA) is initiated. During this process, the maternal transcripts are replaced by newly synthesized and processed RNAs encoded by the embryo’s genome (Jukam et al., 2017).

The nucleolus is the largest sub-nuclear structure of eukaryotic cells where, among other processes, ribosome biogenesis occurs; nucleoli form at so-called nucleolar organizing regions (NORs) and have protein, DNA and RNA components organized in a biomolecular condensate (Lafontaine et al., 2021). As opposed to somatic cells, mammalian oocytes and pre-blastocyst stage embryos contain compact nucleolus precursor bodies (NPBs) of unknown composition. NPBs have been considered to be important for the formation of active nucleoli as sites of aggregation for nucleolar proteins; however, is has been proposed that NPBs may indeed predominantly (or even exclusively) exert their essential function in embryogenesis by establishing chromatin domains; reviewed in (Fulka and Aoki, 2016).

Our objectives were to gain insights into the expression, localization and function of the conserved 3’-5’ exoribonuclease EXOSC10/Rrp6 during embryogenesis in a gene inactivation mouse model.

## Results and Discussion

### Exosc10^−/−^ mice show a prenatal lethality phenotype

We generated a knockout line using *Exosc10*^tm1a(KOMP)Wtsi^ embryonic stem (ES) cells and the “knockout-first” strategy (Austin et al., 2004; Skarnes et al., 2011; Testa et al., 2004) (see Materials and Methods for more details; Fig. 1A). Next, we crossed *Exosc10*^+/−^ male and female mice showing no discernable phenotype to generate offspring. We genotyped 931 newborn mice (including 230 dead newborn pups) and detected 315 wild type mice (33.83%) and 616 heterozygous mice (66.17%) but no homozygous mutants; we note that the *Exosc10*^+/+^ and *Exosc10*^+/−^ alleles show the expected Mendelian distribution (Fig. 1B, C). This result revealed a novel essential role for *Exosc10* during embryo development.

**Figure 1.**
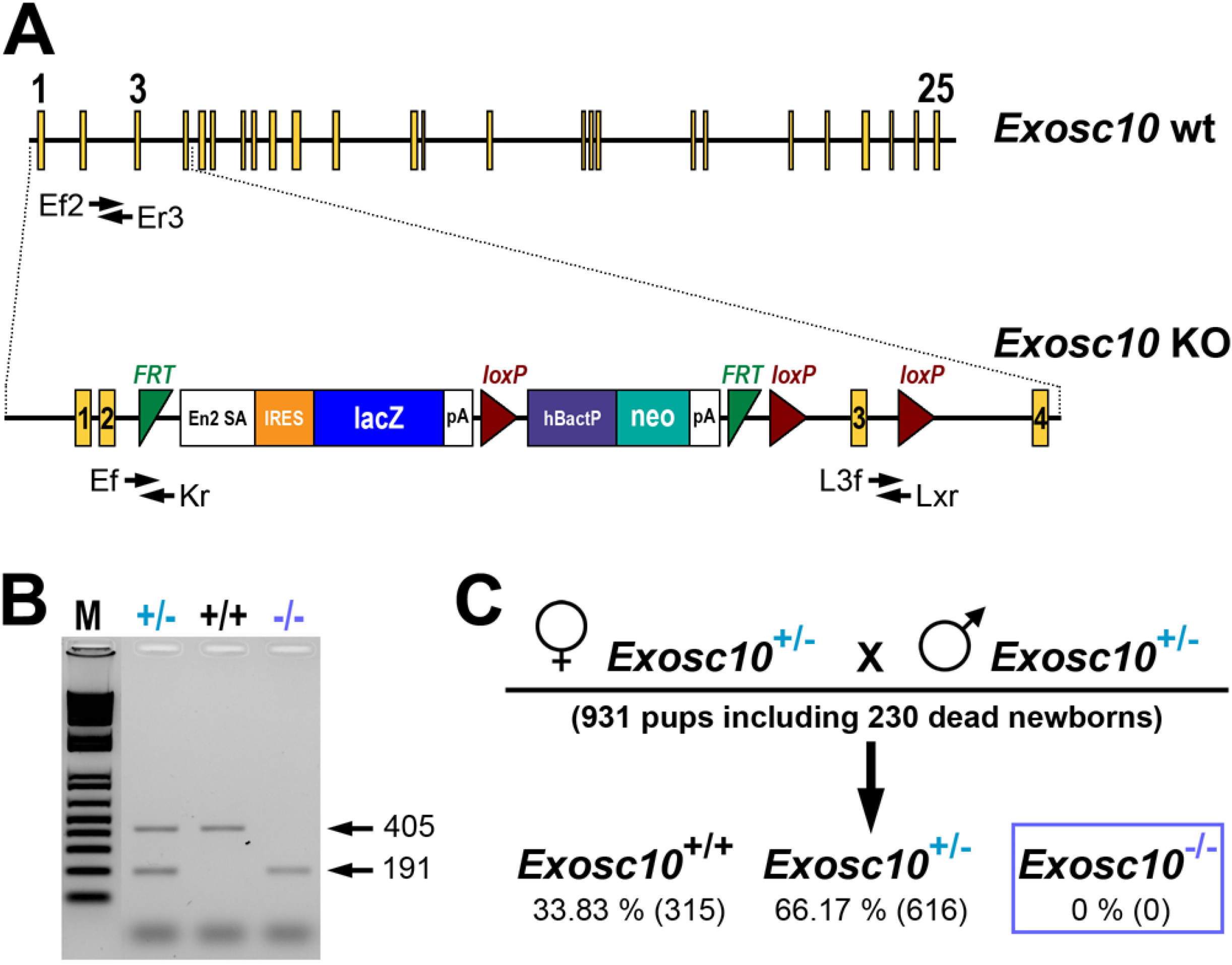
Gene deletion strategy and embryonic lethal phenotype. (A) A schematic shows *Exosc10* wild type (wt) and mutant (KO) alleles. Critical exons are numbered. The knockout-first allele generated by KOMP allows the inactivation of *Exosc10 via* splicing between exon 1-2 and the mouse En2 splice acceptor (En2 SA). Annotated black arrows symbolize PCR primers producing positive amplicons for genotyping and expression assays. (B) A representative genotyping PCR assay is given. Lane 1 contains markers (M) and lanes 2-4 contain samples from wild type (+/+) and heterozygous (+/−) or homozygous mutants (−/−). The PCR amplicon is 405 bp for wt and 191 bp for KO. (C) A chart summarizes the genotypes observed for newborns from heterozygous intercrosses. Percentages are shown and the numbers of mice analysed are given in parentheses.

### Exosc10 is expressed in oocytes and during all stages of early embryogenesis

Based on the results of our genetic analysis we set out to determine the early embryonic expression profile of *Exosc10* and the protein’s subcellular localization pattern in oocytes and blastomeres. Ovarian folliculogenesis is an important process that prepares the mouse oocyte for embryonic development. The oocyte accumulates maternal RNAs and proteins that decline rapidly after fertilization and reach a minimum at the two-cell stage where the major wave of ZGA occurs in the mouse; reviewed in (Li et al., 2010; Schulz and Harrison, 2019; Zhang and Smith, 2015). Successive developmental stages mark the beginning of rRNA gene transcription, the transition from nucleolus precursor bodies (NPB) to mature nucleoli, and embryo compaction when morulae and blastocysts form before the implantation step (Fig. 2A) (Baran et al., 2001; Zatsepina et al., 2003); for review, see (Kresoja-Rakic and Santoro, 2019). Since pre-implantation embryonic development involves maternal and embryonic mRNAs/proteins we sought to gain insights into the expression profile of *Exosc10* during this phase. To this end, we first mated *Exosc10*^+/+^ female mice at the age of two to six months with vasectomized or fertile C57BL/6N males to obtain oocytes and embryos, respectively. Next, we analyzed the expression of *Exosc10* at oocyte and zygote to blastocyst stages by PCR assays using a wild type allele-specific pair of oligonucleotide primers and found that *Exosc10* mRNA is expressed at all stages (Fig. 2B and C). To determine if *Exosc10* mRNA present in the oocyte is of maternal origin we mated *Exosc10*^+/−^ females with vasectomized males, and analysed the oocytes by PCR genotyping for the mutant allele and by RT-PCR for the wild type mRNA. The results show that *Exosc10*^−^ metaphase II (MII) oocytes contain wild type *Exosc10* mRNA (Fig. 2D, lanes 4 and 5).

**Figure 2.**
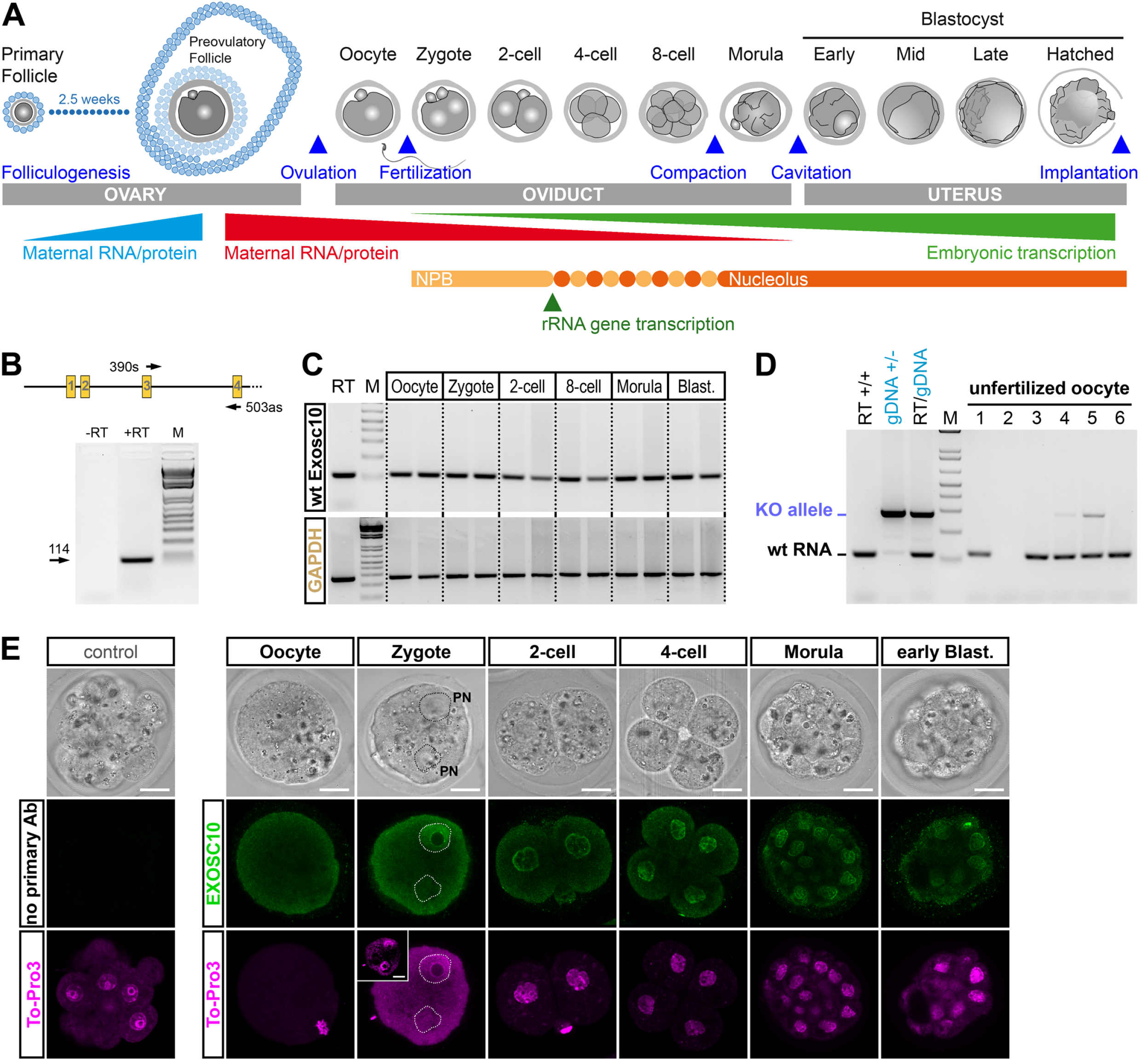
EXOSC10 expression and localization during early embryogenesis. (A) A schematic shows critical phases during folliculogenesis and embryogenesis. Important steps are indicated by blue arrowheads. The timing and levels of maternal and embryonic gene expression patterns, formation of the nucleolus and ribosomal RNA expression are shown. (B) A schematic of the first four *Exosc10* exons locates PCR primers used to assay mRNA expression. The output of PCR expression assays with or without the reverse transcription step (-/+RT) is shown. Lane 3 contains the markers (M). An arrow marks the diagnostic bands for which the size is given in bases. (C) The output of PCR expression assays is shown. Lanes 1 and 2 contain a positive control (RT) and markers (M). Lanes 3-14 contain duplicate samples from an unfertilized egg (Oocyte), a fertilized egg (Zygote), two- and eight-cell embryos (2-cell, 8-cell), morula and blastocyst (Blast). GAPDH was used as a control. (D) The output of PCR expression assays is shown. Lanes 1-4 contain controls for RNA (Testicular RNA, RT+/+), genomic DNA (gDNA+/−), RT/gDNA and markers (M). The remaining lanes contain six samples from female gametes as shown. Diagnostic bands for wild type (wt) and deletion (KO) alleles are indicated. (E) Confocal images of an oocyte, a zygote and early embryos are given. The top row shows brightfield images, the middle and bottom rows show immunofluorescence images of cells assayed for the target protein (EXOSC10) in green and DNA (To-Pro3) in magenta, respectively. The zygote’s pronuclei (PN) are indicated. An insert in the image showing the zygote delineates the localization of the second PN. Scale bars (including the insert) are 20 μm.

### EXOSC10 localizes to the nucleolus precursor body (NPB) in early embryonic cells

In oocytes we observed a diffuse cytoplasmic signal for EXOSC10 (Fig. 2E). In zygotes we detected a diffuse signal in the cytoplasm and the pronuclei (Fig. 2E). From the two-cell stage onward we found that EXOSC10 accumulates in the nucleus (Fig. 2E). In metaphase and anaphase cells, where the nuclear envelope disintegrates and condensed chromosomes are clearly visible, we observe a weak and diffuse cytoplasmic signal (Fig. 3A). However, in newly formed nuclei containing a nuclear envelope and chromatin the protein is concentrated in the nucleus (Fig. 3A, three-cell panels). In four-cell embryos EXOSC10 is also present in foci (Fig. 3B and C).

**Figure 3.**
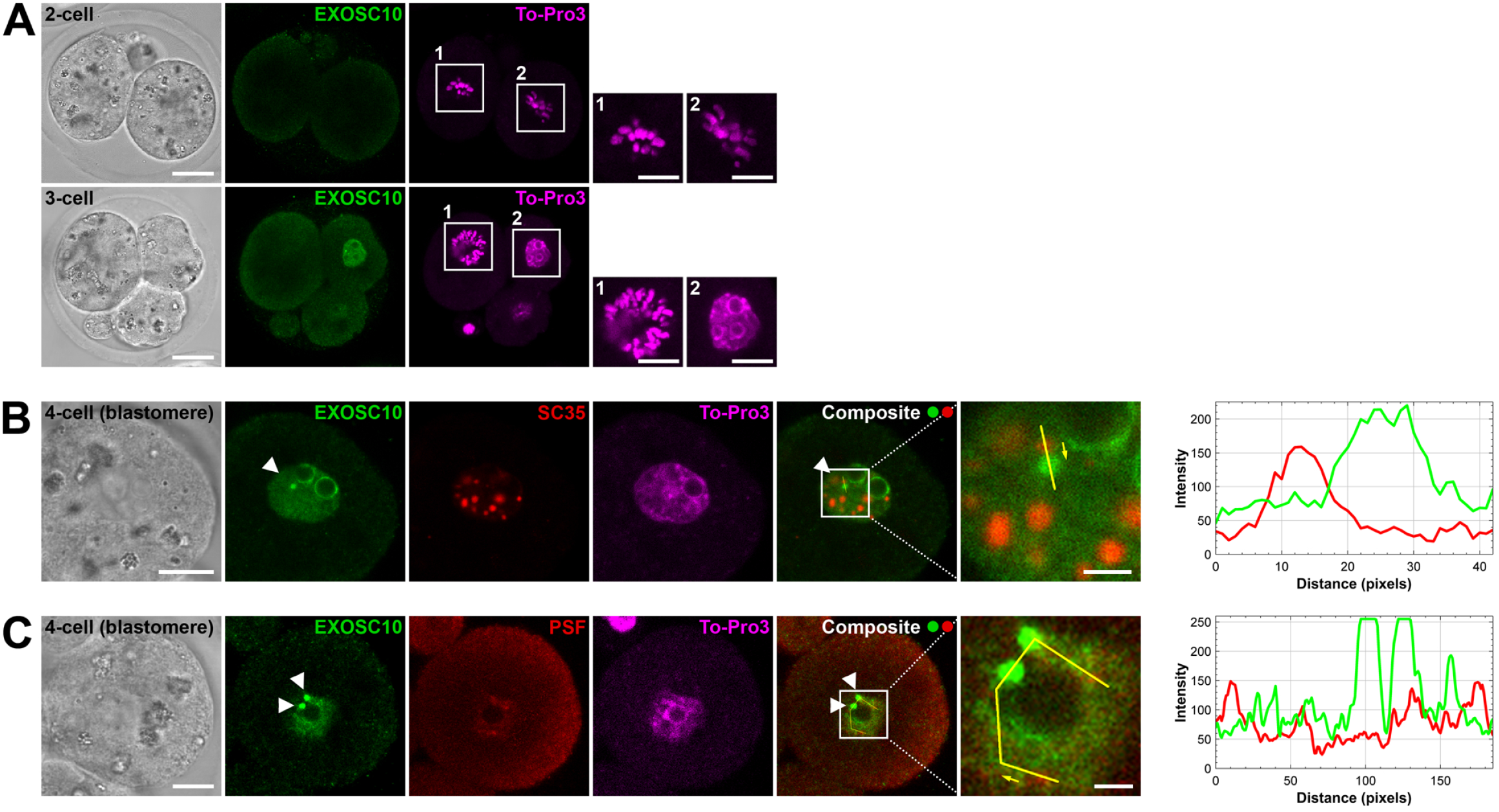
EXOSC10 and subcellular marker protein localization in the early embryo. (A) Confocal images of two- and three-cell embryos as indicated are shown to reveal the morphology (left, light), the target protein (EXOSC10, green) and DNA (To-Pro3, magenta), as indicated. Scale bars are 20 μm and 10 μm for inserts 1 and 2. (B and C) Images of costaining patterns for the target protein (EXOSC10, green), a nuclear speckle marker protein (SC35, red, panel B), a nucleolar cap marker protein (PSF, red, panel C) and DNA (To-Pro3, magenta) in blastomeres from four-cell embryos are shown. Fluorescence intensity profiles of EXOSC10 (green) and SC35/PSF (red) were plotted along the yellow line (yellow arrows indicate the direction of reading) in the composite images. White arrowheads mark EXOSC10-positive foci. Scale bars in B and C are 10 μm and 2.5 μm for the composite inserts.

Eukaryotic cells contain speckles and paraspeckles, which are membraneless structures composed of long non-coding RNAs (lncRNAs), splicing factors and other RNA binding proteins; reviewed in (Chen and Belmont, 2019; Pisani and Baron, 2019). We performed a colocalization assay using marker proteins for speckles (SC35) and paraspeckles (PSF) and found no overlap with the EXOSC10 signal in blastomeres (Fig. 3B and C). This confirms earlier results obtained with cultured fibroblasts and a different anti-EXOSC10 antibody; see Fig. S7 in reference (Fei et al., 2017). Finally, we observed that EXOSC10 accumulates discontinuously at the NPB periphery (Fig. 4A, blastomeres 1a-f and 4g-j) or completely surrounds the NPB, where it colocalizes with the nucleolar marker B23, in different blastomeres of a four- or six-cell embryo (Fig. 4A, blastomeres 2k-l, and Fig. 4B).

**Figure 4.**
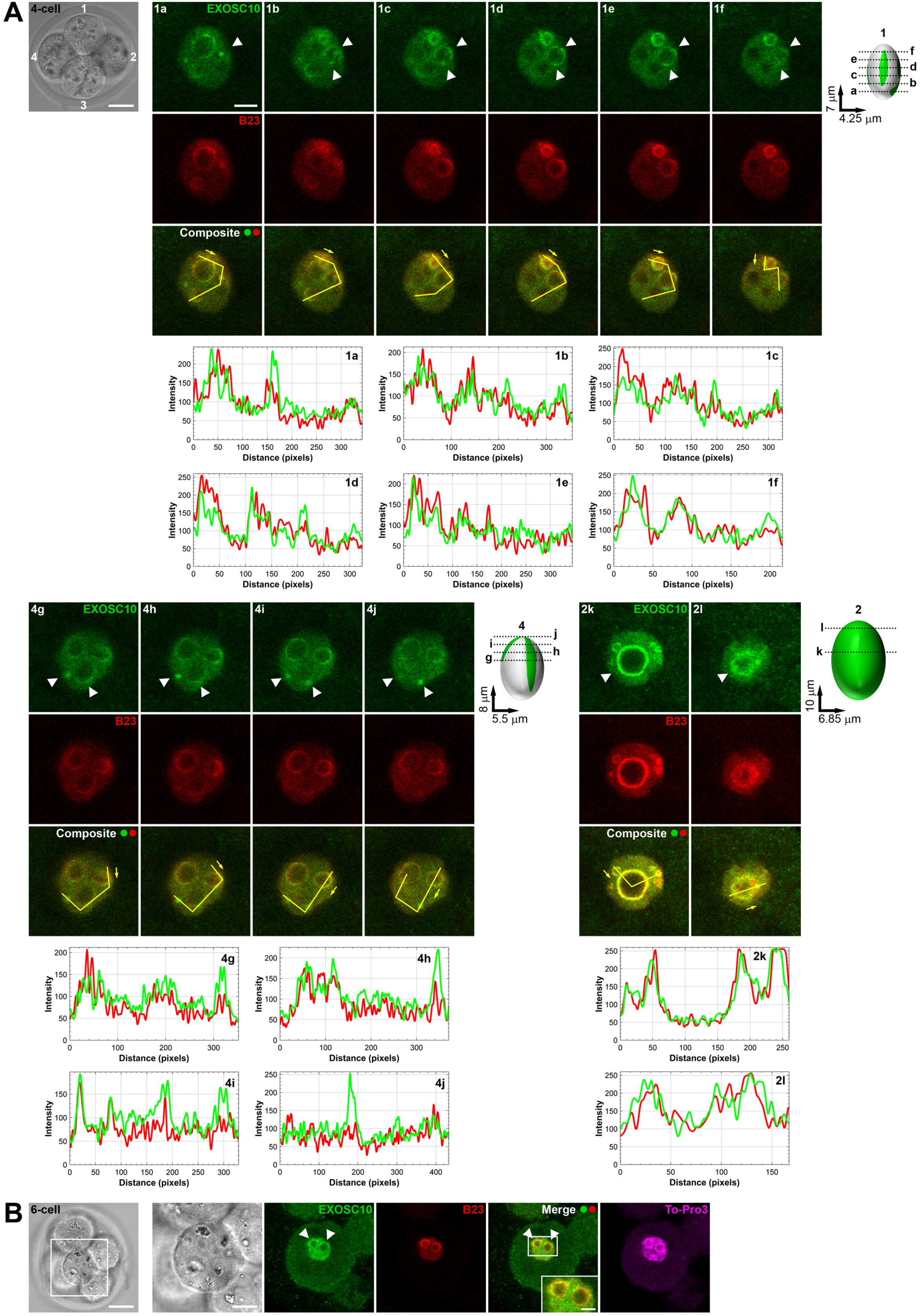
Localization of EXOSC10 at the periphery of the nucleolar precursor bodies. (A) Z-stack images of co-staining patterns for the target protein (EXOSC10, green) and a nucleolar marker protein (B23, red) in a four-cell embryo (brightfield) are shown. Intervals between images are 1 μm for blastomeres 1 and 4, and 3 μm for blastomere 2. Schematics show the positions of confocal sections within individual blastomeres as indicated. The fluorescence intensity was measured along the yellow lines drawn on the composite images (yellow arrows mark the direction of reading) and the corresponding intensity profiles were displayed underneath. White arrowheads mark EXOSC10 foci and partial or complete peripheral signals. Scale bars are 20 μm and 5 μm (1a-f, 4g-j, 2k-l). (B) Confocal images of a six-cell embryo stained for EXOSC10 (green), B23 (red) and DNA (To-Pro3, magenta) are shown. White arrowheads mark EXOSC10 foci and partial or complete peripheral signals. Scale bars are 20 μm for the six-cell embryo brightfield, 10 μm for the enlargement of a blastomere, and 2.5 μm for the merge insert.

Our immunofluorescence data reveal that in blastomeres from the two-cell embryo stage onward EXOSC10 localizes to the periphery of NPBs and to the nucleus, while the protein appears to diffuse into the cytoplasm in oocytes and metaphase blastomeres that lack structured nuclei. Localization to the NPB is in agreement with the protein’s well established function in ribosomal RNA processing (Davidson et al., 2019; Kent et al., 2009) and may point to a potential role in establishing chromatin domains (Chun et al., 2021; Fulka and Aoki, 2016). A critical role for EXOSC10 during cell growth and division via controlling protein translation is also supported by work on the anti-cancer drug 5-FU. This compound inhibits the ribonucleolytic activities of the yeast, fly and human EXOSC10 orthologs, which provides a straightforward explanation for the drug’s interference with late rRNA processing (notably of 5.8S rRNA), ribosome biogenesis and translation elongation by inhibiting the ribonucleolytic activity of EXOSC10 (Abou Elela and Nazar, 1997; Burger et al., 2010; Fang et al., 2004; Hoskins and Scott Butler, 2007; Kammler et al., 2008; Silverstein et al., 2011).

### Exosc10^−/−^ embryos arrest at the eight-cell embryo/morula stage

Next, we asked at which stage homozygous mutant mice undergo a developmental arrest *in utero*. We first analyzed 30 embryos from six independent litters at day e7.5 and observed 10 wild type (33.3%), 20 heterozygous (66.7%), and no homozygous mutant embryos (Fig. 5A). Further analysis of 49 embryos from 10 litters at day e6.5 revealed eight wild type (16.3%), 41 heterozygous (83.7%) and still no homozygous mutant embryos (Fig. 5A). Finally, we genotyped 32 embryos from seven litters at day e4.5 and found 11 wild type (34.4%), 21 heterozygous (65.6%) but no mutant lacking *Exosc10* (Fig. 5A). These results demonstrate a homozygous mutant arrest phenotype during early embryogenesis prior to the embryo implantation step.

**Figure 5.**
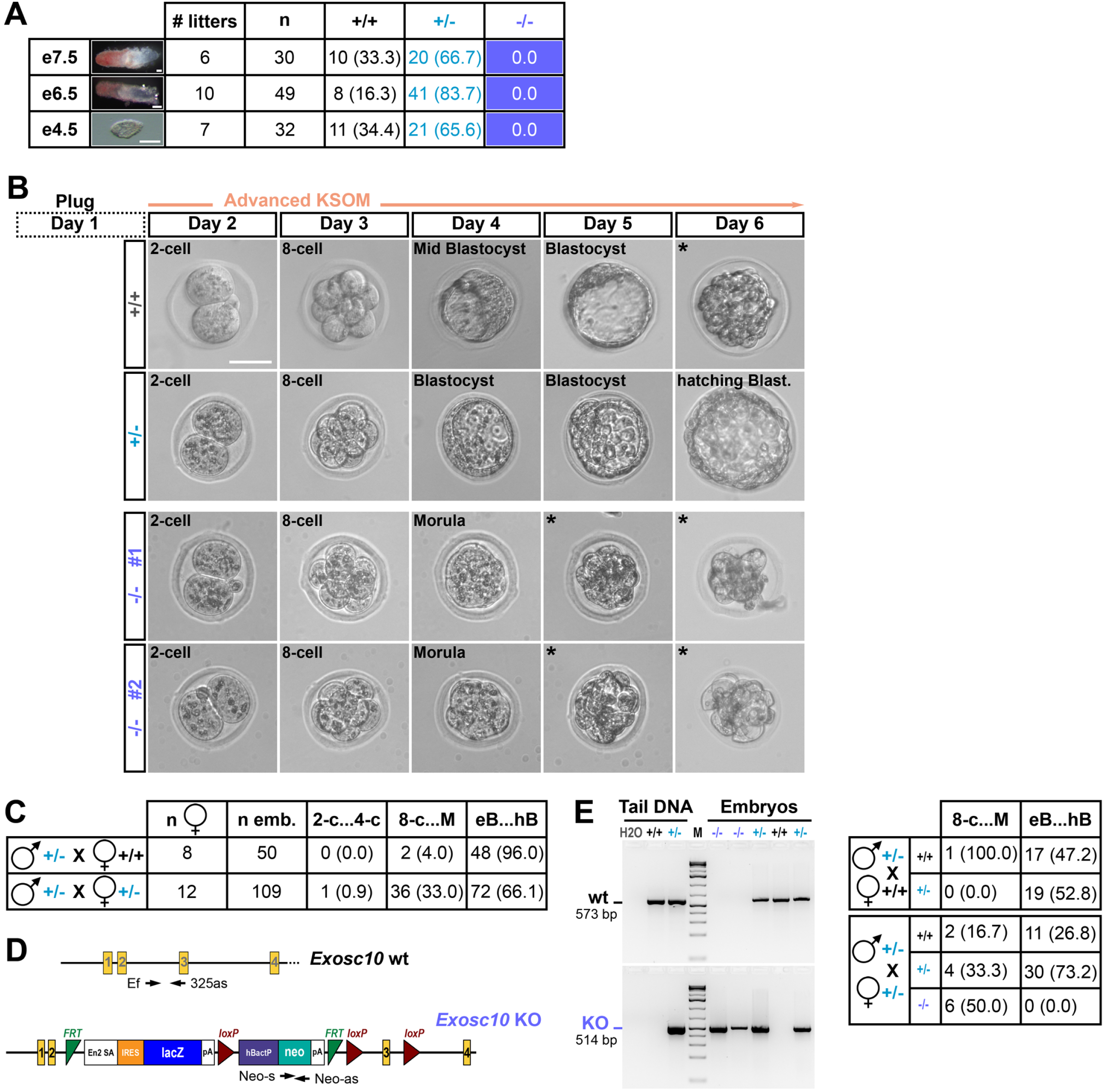
Homozygous mutant developmental arrest phenotype during *in utero* and *in vitro* embryogenesis. (A) A table summarizes the genotypes of embryos collected at e7.5, e6.5 and e4.5 (columns 1 and 2). The numbers of litters (column 3), embryos (column 4) and observed genotypes (columns 5-7) are shown. The data for wild type mice and hetero/homozygous mutants are color-coded for clarity. Numbers of embryos assayed are indicated and percentages are given in parentheses in columns 5 and 6. Scale bars are 100 μm. (B) Representative brightfield images of wild type (+/+), heterozygous (+/−) and homozygous mutant embryos (−/− #1, #2) cultured *in vitro* in Advanced KSOM medium are shown. The initial mating time point (Day 1, Plug) and five additional daily time points (Day2-Day6) are shown at the top. Successive developmental stages are indicated. Black asterisks mark degenerated mutant embryos. Scale bar is 50 μm. (C) A table summarizes the data for embryonic arrest phenotypes determined using offspring from control crosses of heterozygous males and wild type homozygous females (row 1) and experimental intercrosses of heterozygous mice (row 2). Total numbers of females producing litters (n♀) and cultured embryos (n emb), and arrested embryos at the 2- and 4-cell stages (2-c…4-c), 8-cell and morula stages (8-c…M) and early/hatching blastocyst stages (eB…hB) are shown. Percentages are given in parentheses. (D) A schematic shows the location and annotation of PCR primers used to assay wild type (wt) and mutant (KO) *Exosc10* alleles. (E) The output of PCR genotyping assays is shown in the panel to the left. Lanes 1-3 show diagnostic bands obtained with adult control samples as indicated at the top (tail DNA). Lane 4 contains markers (M). Lanes 5-9 show bands obtained with embryo samples showing wild type (+/+), heterozygous (+/−) or homozygous (−/−) genotypes. PCR-positive amplicons are 573 bp for wild type (wt) and 514 bp for mutant (KO) alleles. The table to the right summarizes the genotypes of embryos showing arrest phenotypes at different stages of embryogenesis illustrated in panel B. Numbers of embryos are shown and percentages of the total numbers at different stages are given in parentheses.

Given that maternal wild type *Exosc10* mRNA is present in *Exosc10*^−^ oocytes, we assumed that these embryos are able to reach the two-cell stage before embryonic genome activation occurs and maternal RNA decays. To determine the point at which homozygous mutants arrest beyond that stage, we cultured two-cell embryos for five days. We first asked if our *in vitro* culture conditions allowed for embryogenesis to proceed efficiently (Fig. 5B). To this end, we incubated wild type and heterozygous embryos stemming from crosses between eight *Exosc10*^+/+^ females and *Exosc10*^+/−^ males, and found that our culture conditions did not significantly affect embryo development, since 96% (48/50 embryos) reached the blastocyst stage, while only 4% (2/50) died at the eight-cell embryo/morula stage (Fig. 5C). Next, we analysed embryos stemming from crosses between 12 *Exosc10*^+/−^ females and *Exosc10*^+/−^ males and found that 33% (36/109) undergo a developmental arrest at the eight-cell embryo/morula stage (Fig. 5B and C). Bearing in mind that 25% of the embryos from a heterozygous intercross are expected to be homozygous mutants, and that the vast majority of wild type and heterozygous embryos develop normally, this result suggests that most arrested embryos lack functional EXOSC10.

Finally, we sought to further substantiate our findings by genotyping cultured embryos using wild type and mutant allele-specific primer pairs (Fig. 5D). First, we assayed 37 embryos from control crosses between *Exosc10*^+/+^ females and *Exosc10*^+/−^ males and found one wild type (100%) but no heterozygous mutant at the eight-cell embryo/morula stage, while 17 (47.2%) homozygous and 19 (52.8%) heterozygous embryos developed into blastocysts (Fig. 5E). In comparison, we genotyped 53 embryos from *Exosc10*^+/−^ intercrosses, and observed two wild types (16.7%), four heterozygotes (33.3%) and six homozygotes (50%) among 12 embryos arrested at the eight-cell embryo/morula stage, and 11 wild types (26.8%), 30 heterozygotes (73.2%) and no homozygotes among 41 embryos at the blastocyst stage (Fig. 5E).

Given the known roles of EXOSC10, a plausible interpretation of our data is that the arrest phenotype is due to disrupted nucleolar rRNA processing, which leads to impaired ribosome biogenesis and translation elongation (Abou Elela and Nazar, 1997; Davidson et al., 2019; Knight et al., 2016; Kobylecki et al., 2018). We note that UBF - a co-factor of RNA Polymerase I (POLR1B), which is important for rRNA transcription - shows peripheral NPB localization similar to EXOSC10’s pattern (Kone et al., 2016) and mutant embryos lacking functional UBF or POLR1B arrest at the morula stage (Chen et al., 2008; Hamdane et al., 2017); for review, see (Kresoja-Rakic and Santoro, 2019). Moreover, mice lacking the NPB-associated ribosome biogenesis factor PES1 also show an early phenotype by failing to progress normally through the eight-cell embryo/morula/blastocyst steps (Lerch-Gaggl et al., 2002). In addition to impaired ribosome biogenesis, it is conceivable that altered RNA exosome-dependent and cohesin-mediated genome architecture also contributes to *Exosc10’s* embryonic lethal phenotype (Chun et al., 2021). This is a tempting idea in the context of earlier findings that implicate NPBs and nucleoli in the spatial organization of the genome; reviewed in (Fulka and Aoki, 2016; Kresoja-Rakic and Santoro, 2019).

## Conclusion

Collectively, our results suggest that *Exosc10*^−/−^ mutants arrest at the eight-cell/morula transition prior to formation of the blastocyst. It will be exciting to learn more about the effects disease-related *EXOSC10* alleles might have on cell division, growth and development; reviewed in (Stuparevic et al., 2021), especially since interpreting how mutations might alter protein function is greatly facilitated by the novel AlphaFold protein structure prediction algorithm (Jumper et al., 2021).

## Materials and methods

### Ethics Statement

Studies involving animals, including housing and care, method of euthanasia and experimental protocols were conducted in accordance with the French regulation for Laboratory Animal Care, issued from the transposition of the European Directive 2010/63/UE. This Directive stipulates in article 1, alinea 2 that euthanizing laboratory animals for the sole purpose of investigating their tissues does not require prior authorization by an ethics committee. Vasectomized mice used in this study were part of a project approved by the French ministry of research (APAFIS#25237-2020042715505401 v2). The animal facility was licensed by the French Ministry of Agriculture (C35-238-19). Animals were sacrificed using CO_2_, according to a method in Annex IV of the directive. All experiments were supervised by Fabrice G. Petit who is licensed for animal experimentation by the French Ministry of Agriculture (A92–313).

### *Generation* of *Exosc10* knockout (KO) mice

We generated the *Exosc10* KO mouse strain using *Exosc10*^tm1a(KOMP)Wtsi^ embryonic stem (ES) cell clones EPD0567_5_B04 and EPD0567_5_F03, which were provided by the Knockout Mouse Project (KOMP) funded by the NIH and obtained *via* the KOMP Repository (www.komp.org) funded by the NCRR-NIH (Jamin et al., 2017). *Exosc10* knockout first, promoter driven mice were generated based on the “knockout-first” design, where a trapping cassette including the *En2* splice acceptor (En2 SA) and SV40 polyadenylation (pA) sequences is inserted within the second intron of *Exosc10* such that splicing between the first and second exons and En2 SA creates a knockout *Exosc10* allele (Skarnes et al., 2011; Testa et al., 2004). ES cell clones were used to generate chimeric mice at Institut Clinique de la Souris (ICS, www.ics-mci.fr). After germline transmission, *Exosc10*^+/−^ mice were transferred to our local animal facility.

### Two-cell embryo culture

We mated *Exosc10*^+/+^ or *Exosc10*^+/−^ females with stud *Exosc10*^+/−^ male mice and recovered two-cell stage (gestational day 2; GD2) embryos by flushing approximately 0.5 ml of EmbryoMax Advanced KSOM medium (Merck-Sigma, MR-101-D) into the oviduct. We placed embryos in a 96-well plate containing 100 μl of Advanced KSOM medium, cultured them in a humidified incubator with 5% CO_2_ at 37°C and monitored them daily (2-4PM) from day two (two-cell stage, GD2) to day six. Embryos where collected and rinsed three times in H_2_O before processing them for PCR genotyping. We documented the process using a CELENA S Digital Imaging System (Logos Biosystems).

### Unfertilized oocyte and embryo recovery

We obtained unfertilized oocytes by mating *Exosc10*^+/+^ or *Exosc10*^+/−^ female mice with vasectomized males. Mice were sacrificed the day a vaginal plug was observed (GD1) or e0.5). Under a stereo dissecting microscope we cut the ampulla region of the oviduct and collected the cumulus oocyte mass in prewarmed M2 medium (Sigma, France). We removed the cumulus mass from ovulated eggs by incubating them for 5-10 min in M2 medium supplemented with 1 mg/ml hyaluronidase (Sigma) at 37°C. To recover embryos from zygote to morula stages (GD1 to GD3), we inserted a 30-gauge needle attached to a 1 ml syringe filled with prewarmed M2 into the oviduct and flushed the embryos out. To collect blastocysts and post implantation embryos (GD4 and GD6 to GD8), we inserted a 27-gauge needle attached to a 1 ml syringe filled with M2 into the uterine horn and flushed the embryos out. Oocytes and embryos were rinsed twice in M2 and three times in H_2_O before processing for reverse transcription or PCR.

### Oligonucleotide primers for mouse and embryo genotyping and the detection of mRNA

Ef2 – TCT AAG CCT GAC AGC ATG AAT TGA ACC

Er3 – GTG GTG TAG ACT TGT GAT ACT GC

L3f – TGC AAG GAG AAA CCC CAG TGT TAC G

Lxr – TTA TCA TTA ATT GCG TTG CGC CAT C

Ef – GAT GGA GCG AGC AAG CTT CTG

Kr – CCA ACA GCT TCC CCA CAA CGG

325as – CTG CGA CAG CCA TGG TAC T

390s – AAT GAC GTG ATA TTG GAG AGA GTG

503as – GGA TAC TAT TGT TTT GGG GAC CT

Neo-s – GCC CTG AAT GAA CTG CAG G

Neo-as – CAC GGG TAG CCA ACG CTA T

GAPDH-s – TGG ATC TGA CAT GCC GCC TG

GAPDH-as – ATG TAG GCC ATG AGG TCC AC

### Mouse genotyping

We digested tail tips collected at weaning and from sacrificed animals in 600 μl of 50 mM NaOH at 95°C for 60 min and before adding 50 μl of 1 M Tris-HCl pH 8.0 and centrifuging samples as 13,400 x g for 5 min. Then we added 1 μl of supernatant to a final volume of 15 μl containing 1x CoralLoad PCR buffer, 0.1 mM dNTP (Invitrogen, France), 0.1 mM mixed primers (Ef2/Er3/L3f/Lxr, 1:1:1:1, Eurogentec, France), 1x Q solution and 0.017 units of HotStar Taq Plus DNA polymerase (Qiagen, France). PCR was at 95°C for 5 min, 35 cycles at 94°C for 1 min, 58°C for 1 min, 72°C for 1 min and 72°C for 10 min.

### Embryo genotyping

We collected embryos at e4.5, e6.5 and e7.5 and digested them in 6 μl of 50 mM NaOH at 95°C for 10 min before adding 0.5 μl of 1 M Tris-HCl pH 8.0. For genotyping, we set the parameter at 40 cycles. For cultured embryos, we lysed samples in 8 μl of lysis buffer PBND containing 0.1 mg/ml Proteinase K (Roche) (Scavizzi et al., 2015) and incubated them at 56°C for 30 min before we inactivated Proteinase K at 95°C for 10 min. PCR was set at 45 cycles using Ef/325as for wild type and Neo-s/Neo-as for mutant alleles.

### Reverse transcription and PCR analysis

We isolated total RNA from oocytes to blastocysts using the Arcturus PicoPure kit (Applied Biosystems). We extracted RNA in 50 μl of Extraction buffer, eluted total RNA in 10 μl of Elution buffer and reverse transcribed RNA in 20 μl using the iScript cDNA synthesis kit (Bio-Rad). We incubated the samples for 5 min at 25°C, 20 min at 46°C and 1 min at 95°C. PCR was performed using 2 μl of RT sample following the mouse genotyping protocol for 42 cycles. We used 390s/503as primers for wild type *Exosc10* mRNA and GAPDH-s/GAPDH-as primers as a control.

### Detection of the maternal Exosc10 mRNA

We obtained unfertilized MII oocytes from *Exosc10*^+/−^ female mice and rinsed them in H_2_O before transferring them into a PCR tube in a variable volume of 1-5 μl H_2_O. We processed samples in a thermal cycler during three cycles at 65°C for 5 min and 4°C for 1 min. Subsequently, the RNA was reverse transcribed (RT) using the iScript cDNA synthesis kit (Bio-Rad) in a final volume of 10 μl. Therefore, the resulting samples contain both genomic DNA and cDNA. We assayed the KO allele and the wt mRNA using Ef/Kr and 390s/503as primer sets, respectively. We performed PCRs at 45 cycles using 10 μl of RT sample in a final volume of 20 μl.

### Immunofluorescence (IF) staining

For IF/confocal analyses 27 *Exosc10*^+/+^ or C57BL/6NRj female mice (Janvier Labs, Le Genest Saint Isle, France) were used to collect 52 oocytes, 35 zygotes, 46 two-cell embryos, 83 four-cell embryos, seven six/eight-cell embryos, eight morulae and one early blastocyst. Confocal images were generated with four oocytes, six zygotes, 12 two-cell embryos, 23 four-cell embryos, three six/eight-cell embryos, four morulae and one early blastocyst. We collected unfertilized oocytes and embryos in M2 medium and fixed them in 4% paraformaldehyde (Electron Microscopy Sciences) 1x PBS for 30 min at room temperature. We permeabilized the samples in 5% BSA-0.5% Triton X-100 1x PBS for 30 min at room temperature before rinsing them in blocking buffer (5% BSA in 1x PBS). The first antibody diluted in blocking buffer (1:200) was incubated at 4°C overnight. A few samples were incubated in blocking buffer only (“no primary antibody”) as a negative control. The next day, we washed the samples three times in 1x PBS before adding the secondary antibody (1:500 in blocking buffer) for 60 min at room temperature. We washed the samples three times in 1x PBS and kept them in a 1% BSA 1x PBS solution until mounting. Antibodies used in this study were against EXOSC10 (Abcam, ab50558), B23 (Sigma, B0556), PSF (Sigma, P2860) and SC-35 (Sigma, S4045). Secondary antibodies were Alexa Fluor 488 (Life Technologies, chicken anti-rabbit antibody A-21441) and Alexa Fluor 555 (Life Technologies, donkey anti-mouse antibody A-31570).

### Confocal imaging, image processing and analysis

We kept unfertilized oocytes and embryos (zygotes to blastocysts) in 1% BSA 1x PBS after immunofluorescence staining before we placed them on glass-bottom dishes (MatTek Life Sciences, Ashland, MA, USA) for inspection with a Leica SP5 confocal microscope equipped with a Plan-Apo 63x/1.4 HC oil immersion objective and lasers with excitation lines at 488 nm, 561 nm and 633 nm. We used To-Pro-3 to stain DNA (far red-fluorescent stain, Invitrogen). We acquired image stacks at 1 μm intervals and processed them using the Application Suite Advanced Fluorescence software (Leica). ImageJ (version 1.53j) software available from the NIH (https://imagej.nih.gov/ij/) was used to create composite images by merging the red and green channels and to obtain intensity RGB profile plots.

## Acknowledgments

We thank the Knockout Mouse Project (KOMP), the Institut Clinique de la Souris (ICS) and the UMS 3480 – US 018 Biosit Facility Ibisa Microscopy Rennes Imaging Center (MRic-Photonics) for providing excellent services.

## References

Abou Elela, S., Nazar, R.N., 1997. Role of the 5.8S rRNA in ribosome translocation. Nucleic Acids Res. 25, 1788–1794.

Austin, C.P., Battey, J.F., Bradley, A., Bucan, M., Capecchi, M., Collins, F.S., Dove, W.F., Duyk, G., Dymecki, S., Eppig, J.T., Grieder, F.B., Heintz, N., Hicks, G., Insel, T.R., Joyner, A., Koller, B.H., Lloyd, K.C., Magnuson, T., Moore, M.W., Nagy, A., Pollock, J.D., Roses, A.D., Sands, A.T., Seed, B., Skarnes, W.C., Snoddy, J., Soriano, P., Stewart, D.J., Stewart, F., Stillman, B., Varmus, H., Varticovski, L., Verma, I.M., Vogt, T.F., von Melchner, H., Witkowski, J., Woychik, R.P., Wurst, W., Yancopoulos, G.D., Young, S.G., Zambrowicz, B., 2004. The knockout mouse project. Nat. Genet. 36, 921–924.

Baran, V., Brochard, V., Renard, J.P., Flechon, J.E., 2001. Nopp 140 involvement in nucleologenesis of mouse preimplantation embryos. Mol. Reprod. Dev. 59, 277–284.

Blomen, V.A., Majek, P., Jae, L.T., Bigenzahn, J.W., Nieuwenhuis, J., Staring, J., Sacco, R., van Diemen, F.R., Olk, N., Stukalov, A., Marceau, C., Janssen, H., Carette, J.E., Bennett, K.L., Colinge, J., Superti-Furga, G., Brummelkamp, T.R., 2015. Gene essentiality and synthetic lethality in haploid human cells. Science 350, 1092–1096.

Bluthner, M., Bautz, F.A., 1992. Cloning and characterization of the cDNA coding for a polymyositis-scleroderma overlap syndrome-related nucleolar 100-kD protein. J. Exp. Med. 176, 973–980.

Briggs, M.W., Burkard, K.T., Butler, J.S., 1998. Rrp6p, the yeast homologue of the human PM-Scl 100-kDa autoantigen, is essential for efficient 5.8 S rRNA 3’ end formation. J. Biol. Chem. 273, 13255–13263.

Burger, K., Muhl, B., Harasim, T., Rohrmoser, M., Malamoussi, A., Orban, M., Kellner, M., Gruber-Eber, A., Kremmer, E., Holzel, M., Eick, D., 2010. Chemotherapeutic drugs inhibit ribosome biogenesis at various levels. J. Biol. Chem. 285, 12416–12425.

Chazaud, C., Yamanaka, Y., 2016. Lineage specification in the mouse preimplantation embryo. Development 143, 1063–1074.

Chen, H., Li, Z., Haruna, K., Li, Z., Li, Z., Semba, K., Araki, M., Yamamura, K., Araki, K., 2008. Early pre-implantation lethality in mice carrying truncated mutation in the RNA polymerase 1-2 gene. Biochem. Biophys. Res. Commun. 365, 636–642.

Chen, Y., Belmont, A.S., 2019. Genome organization around nuclear speckles. Curr. Opin. Genet. Dev. 55, 91–99.

Chun, Y., Han, S., Kim, T., Cho, Y., Lee, D., 2021. RNA surveillance controls 3D genome structure via stable cohesin-chromosome interaction. bioRxiv.

Clift, D., Schuh, M., 2013. Restarting life: fertilization and the transition from meiosis to mitosis. Nat. Rev. Mol. Cell Biol. 14, 549–562.

Davidson, L., Francis, L., Cordiner, R.A., Eaton, J.D., Estell, C., Macias, S., Caceres, J.F., West, S., 2019. Rapid Depletion of DIS3, EXOSC10, or XRN2 Reveals the Immediate Impact of Exoribonucleolysis on Nuclear RNA Metabolism and Transcriptional Control. Cell Rep. 26, 2779–2791 e2775.

Domingo-Prim, J., Endara-Coll, M., Bonath, F., Jimeno, S., Prados-Carvajal, R., Friedlander, M.R., Huertas, P., Visa, N., 2019. EXOSC10 is required for RPA assembly and controlled DNA end resection at DNA double-strand breaks. Nat Commun 10, 2135.

Eberle, A.B., Jordan-Pla, A., Ganez-Zapater, A., Hessle, V., Silberberg, G., von Euler, A., Silverstein, R.A., Visa, N., 2015. An Interaction between RRP6 and SU(VAR)3-9 Targets RRP6 to Heterochromatin and Contributes to Heterochromatin Maintenance in Drosophila melanogaster. PLoS Genet. 11, e1005523.

Fang, F., Hoskins, J., Butler, J.S., 2004. 5-fluorouracil enhances exosome-dependent accumulation of polyadenylated rRNAs. Mol. Cell. Biol. 24, 10766–10776.

Fei, J., Jadaliha, M., Harmon, T.S., Li, I.T.S., Hua, B., Hao, Q., Holehouse, A.S., Reyer, M., Sun, Q., Freier, S.M., Pappu, R.V., Prasanth, K.V., Ha, T., 2017. Quantitative analysis of multilayer organization of proteins and RNA in nuclear speckles at super resolution. J. Cell Sci. 130, 4180–4192.

Fulka, H., Aoki, F., 2016. Nucleolus Precursor Bodies and Ribosome Biogenesis in Early Mammalian Embryos: Old Theories and New Discoveries. Biol. Reprod. 94, 143.

Ge, Q., Frank, M.B., O’Brien, C., Targoff, I.N., 1992. Cloning of a complementary DNA coding for the 100-kD antigenic protein of the PM-Scl autoantigen. J. Clin. Invest. 90, 559–570.

Gudipati, R.K., Xu, Z., Lebreton, A., Seraphin, B., Steinmetz, L.M., Jacquier, A., Libri, D., 2012. Extensive degradation of RNA precursors by the exosome in wild-type cells. Mol. Cell 48, 409–421.

Hamdane, N., Tremblay, M.G., Dillinger, S., Stefanovsky, V.Y., Nemeth, A., Moss, T., 2017. Disruption of the UBF gene induces aberrant somatic nucleolar bodies and disrupts embryo nucleolar precursor bodies. Gene 612, 5–11.

Hoskins, J., Scott Butler, J., 2007. Evidence for distinct DNA- and RNA-based mechanisms of 5-fluorouracil cytotoxicity in Saccharomyces cerevisiae. Yeast 24, 861–870.

Jamin, S.P., Petit, F.G., Kervarrec, C., Smagulova, F., Illner, D., Scherthan, H., Primig, M., 2017. EXOSC10/Rrp6 is post-translationally regulated in male germ cells and controls the onset of spermatogenesis. Sci. Rep. 7, 15065.

Jukam, D., Shariati, S.A.M., Skotheim, J.M., 2017. Zygotic Genome Activation in Vertebrates. Dev. Cell 42, 316–332.

Jumper, J., Evans, R., Pritzel, A., Green, T., Figurnov, M., Ronneberger, O., Tunyasuvunakool, K., Bates, R., Zidek, A., Potapenko, A., Bridgland, A., Meyer, C., Kohl, S.A.A., Ballard, A.J., Cowie, A., Romera-Paredes, B., Nikolov, S., Jain, R., Adler, J., Back, T., Petersen, S., Reiman, D., Clancy, E., Zielinski, M., Steinegger, M., Pacholska, M., Berghammer, T., Bodenstein, S., Silver, D., Vinyals, O., Senior, A.W., Kavukcuoglu, K., Kohli, P., Hassabis, D., 2021. Highly accurate protein structure prediction with AlphaFold. Nature 596, 583–589.

Kammler, S., Lykke-Andersen, S., Jensen, T.H., 2008. The RNA exosome component hRrp6 is a target for 5-fluorouracil in human cells. Mol. Cancer Res. 6, 990–995.

Kent, T., Lapik, Y.R., Pestov, D.G., 2009. The 5’ external transcribed spacer in mouse ribosomal RNA contains two cleavage sites. RNA 15, 14–20.

Kilchert, C., Wittmann, S., Vasiljeva, L., 2016. The regulation and functions of the nuclear RNA exosome complex. Nat. Rev. Mol. Cell Biol. 17, 227–239.

Knight, J.R., Bastide, A., Peretti, D., Roobol, A., Roobol, J., Mallucci, G.R., Smales, C.M., Willis, A.E., 2016. Cooling-induced SUMOylation of EXOSC10 down-regulates ribosome biogenesis. RNA 22, 623–635.

Kobylecki, K., Drazkowska, K., Kulinski, T.M., Dziembowski, A., Tomecki, R., 2018. Elimination of 01/A’-A0 pre-rRNA processing by-product in human cells involves cooperative action of two nuclear exosome-associated nucleases: RRP6 and DIS3. RNA 24, 1677–1692.

Kone, M.C., Fleurot, R., Chebrout, M., Debey, P., Beaujean, N., Bonnet-Garnier, A., 2016. Three-Dimensional Distribution of UBF and Nopp140 in Relationship to Ribosomal DNA Transcription During Mouse Preimplantation Development. Biol. Reprod. 94, 95.

Kresoja-Rakic, J., Santoro, R., 2019. Nucleolus and rRNA Gene Chromatin in Early Embryo Development. Trends Genet. 35, 868–879.

Kuai, L., Das, B., Sherman, F., 2005. A nuclear degradation pathway controls the abundance of normal mRNAs in Saccharomyces cerevisiae. Proc. Natl. Acad. Sci. U. S. A. 102, 13962–13967.

Lafontaine, D.L.J., Riback, J.A., Bascetin, R., Brangwynne, C.P., 2021. The nucleolus as a multiphase liquid condensate. Nat. Rev. Mol. Cell Biol. 22, 165–182.

Lardenois, A., Liu, Y., Walther, T., Chalmel, F., Evrard, B., Granovskaia, M., Chu, A., Davis, R.W., Steinmetz, L.M., Primig, M., 2011. Execution of the meiotic noncoding RNA expression program and the onset of gametogenesis in yeast require the conserved exosome subunit Rrp6. Proc. Natl. Acad. Sci. U. S. A. 108, 1058–1063.

Lerch-Gaggl, A., Haque, J., Li, J., Ning, G., Traktman, P., Duncan, S.A., 2002. Pescadillo is essential for nucleolar assembly, ribosome biogenesis, and mammalian cell proliferation. J. Biol. Chem. 277, 45347–45355.

Li, L., Zheng, P., Dean, J., 2010. Maternal control of early mouse development. Development 137, 859–870.

Mahler, M., Raijmakers, R., 2007. Novel aspects of autoantibodies to the PM/Scl complex: clinical, genetic and diagnostic insights. Autoimmun. Rev. 6, 432–437.

Makino, D.L., Baumgartner, M., Conti, E., 2013. Crystal structure of an RNA-bound 11-subunit eukaryotic exosome complex. Nature 495, 70–75.

Marin-Vicente, C., Domingo-Prim, J., Eberle, A.B., Visa, N., 2015. RRP6/EXOSC10 is required for the repair of DNA double-strand breaks by homologous recombination. J. Cell Sci. 128, 1097–1107.

Mehta, C., Fraga de Andrade, I., Matson, D.R., Dewey, C.N., Bresnick, E.H., 2021. RNA-regulatory exosome complex confers cellular survival to promote erythropoiesis. Nucleic Acids Res.

Meldal, B.H., Forner-Martinez, O., Costanzo, M.C., Dana, J., Demeter, J., Dumousseau, M., Dwight, S.S., Gaulton, A., Licata, L., Melidoni, A.N., Ricard-Blum, S., Roechert, B., Skyzypek, M.S., Tiwari, M., Velankar, S., Wong, E.D., Hermjakob, H., Orchard, S., 2015. The complex portal--an encyclopaedia of macromolecular complexes. Nucleic Acids Res. 43, D479–484.

Midtgaard, S.F., Assenholt, J., Jonstrup, A.T., Van, L.B., Jensen, T.H., Brodersen, D.E., 2006. Structure of the nuclear exosome component Rrp6p reveals an interplay between the active site and the HRDC domain. Proc. Natl. Acad. Sci. U. S. A. 103, 11898–11903.

Neil, H., Malabat, C., d’Aubenton-Carafa, Y., Xu, Z., Steinmetz, L.M., Jacquier, A., 2009. Widespread bidirectional promoters are the major source of cryptic transcripts in yeast. Nature 457, 1038–1042.

Pefanis, E., Wang, J., Rothschild, G., Lim, J., Kazadi, D., Sun, J., Federation, A., Chao, J., Elliott, O., Liu, Z.P., Economides, A.N., Bradner, J.E., Rabadan, R., Basu, U., 2015. RNA exosome-regulated long non-coding RNA transcription controls super-enhancer activity. Cell 161, 774–789.

Pisani, G., Baron, B., 2019. Nuclear paraspeckles function in mediating gene regulatory and apoptotic pathways. Noncoding RNA Res 4, 128–134.

Scavizzi, F., Ryder, E., Newman, S., Raspa, M., Gleeson, D., Wardle-Jones, H., Montoliu, L., Fernandez, A., Dessain, M.L., Larrigaldie, V., Khorshidi, Z., Vuolteenaho, R., Soininen, R., Andre, P., Jacquot, S., Hong, Y., de Angelis, M.H., Ramirez-Solis, R., Doe, B., 2015. Blastocyst genotyping for quality control of mouse mutant archives: an ethical and economical approach. Transgenic Res. 24, 921–927.

Schneider, C., Kudla, G., Wlotzka, W., Tuck, A., Tollervey, D., 2012. Transcriptome-wide analysis of exosome targets. Mol. Cell 48, 422–433.

Schulz, K.N., Harrison, M.M., 2019. Mechanisms regulating zygotic genome activation. Nat. Rev. Genet. 20, 221–234.

Silverstein, R.A., Gonzalez de Valdivia, E., Visa, N., 2011. The incorporation of 5-fluorouracil into RNA affects the ribonucleolytic activity of the exosome subunit Rrp6. Mol. Cancer Res. 9, 332–340.

Skarnes, W.C., Rosen, B., West, A.P., Koutsourakis, M., Bushell, W., Iyer, V., Mujica, A.O., Thomas, M., Harrow, J., Cox, T., Jackson, D., Severin, J., Biggs, P., Fu, J., Nefedov, M., de Jong, P.J., Stewart, A.F., Bradley, A., 2011. A conditional knockout resource for the genomewide study of mouse gene function. Nature 474, 337–342.

Stuparevic, I., Novacic, A., Rahmouni, A.R., Fernandez, A., Lamb, N., Primig, M., 2021. Regulation of the conserved 3’-5’ exoribonuclease EXOSC10/Rrp6 during cell division, development and cancer. Biol. Rev. Camb. Philos. Soc.

Testa, G., Schaft, J., van der Hoeven, F., Glaser, S., Anastassiadis, K., Zhang, Y., Hermann, T., Stremmel, W., Stewart, A.F., 2004. A reliable lacZ expression reporter cassette for multipurpose, knockout-first alleles. Genesis 38, 151–158.

Ulmke, P.A., Xie, Y., Sokpor, G., Pham, L., Shomroni, O., Berulava, T., Rosenbusch, J., Basu, U., Fischer, A., Nguyen, H.P., Staiger, J.F., Tuoc, T., 2021. Post-transcriptional regulation by the exosome complex is required for cell survival and forebrain development via repression of P53 signaling. Development 148.

Wagschal, A., Rousset, E., Basavarajaiah, P., Contreras, X., Harwig, A., Laurent-Chabalier, S., Nakamura, M., Chen, X., Zhang, K., Meziane, O., Boyer, F., Parrinello, H., Berkhout, B., Terzian, C., Benkirane, M., Kiernan, R., 2012. Microprocessor, Setx, Xrn2, and Rrp6 cooperate to induce premature termination of transcription by RNAPII. Cell 150, 1147–1157.

Wasmuth, E.V., Januszyk, K., Lima, C.D., 2014. Structure of an Rrp6-RNA exosome complex bound to poly(A) RNA. Nature 511, 435–439.

Wasmuth, E.V., Lima, C.D., 2017. The Rrp6 C-terminal domain binds RNA and activates the nuclear RNA exosome. Nucleic Acids Res. 45, 846–860.

Wu, D., Dean, J., 2020. EXOSC10 sculpts the transcriptome during the growth-to-maturation transition in mouse oocytes. Nucleic Acids Res. 48, 5349–5365.

Xu, Z., Wei, W., Gagneur, J., Perocchi, F., Clauder-Munster, S., Camblong, J., Guffanti, E., Stutz, F., Huber, W., Steinmetz, L.M., 2009. Bidirectional promoters generate pervasive transcription in yeast. Nature 457, 1033–1037.

Zatsepina, O., Baly, C., Chebrout, M., Debey, P., 2003. The step-wise assembly of a functional nucleolus in preimplantation mouse embryos involves the cajal (coiled) body. Dev. Biol. 253, 66–83.

Zernicka-Goetz, M., Morris, S.A., Bruce, A.W., 2009. Making a firm decision: multifaceted regulation of cell fate in the early mouse embryo. Nat. Rev. Genet. 10, 467–477.

Zhang, K., Smith, G.W., 2015. Maternal control of early embryogenesis in mammals. Reprod. Fertil. Dev. 27, 880–896.

